# Passive Acoustic Monitoring within the Northwest Forest Plan Area: 2024 Annual Report

**DOI:** 10.64898/2025.12.18.695274

**Authors:** Damon B. Lesmeister, Julianna M. A. Jenkins, Zachary J. Ruff, Natalie M. Rugg, Simeon Abidari, Sallie Beavers, Roger Christophersen, Rita Claremont, Raymond J. Davis, Scott Gremel, Aaron Henderson, Brandon Henson, Julia Kasper, Christopher McCafferty, Alejandra Muñoz, Dave Press, Courtney Quinn, Suzanne Reffler, James K. Swingle, Erica Tevini, Alaina D. Thomas, Kira Ware, Kirsten Wert

## Abstract

Passive acoustic monitoring (PAM) has become the primary framework for assessing the status, distribution, and habitat associations of forest-dependent wildlife under the Northwest Forest Plan (NWFP). In 2024, the NWFP PAM program continued to operate as a fully integrated, range-wide monitoring network, building on deployments initiated in 2018 and now conducting surveys across approximately 24 million acres of federal land. Using a randomized 2+20% sampling design, more than 4,000 autonomous recording units generated over 2 million hours of acoustic data analyzed with convolutional neural networks, enabling statistically rigorous inference at regional and range-wide scales. This report summarizes 2024 field efforts and analytical advancements in the context of recent peer-reviewed and in-progress studies. Updated analyses confirm persistent, low occupancy of northern spotted owls (*Strix occidentalis caurina*), with detections concentrated in the Klamath Mountains and southern Cascades, consistent with long-term demographic declines. Range-wide models of barred owl (*Strix varia*) landscape use continue to show high occupancy across much of the NWFP area, reinforcing competitive displacement as the dominant driver of spotted owl range contraction. The PAM network also advanced monitoring of marbled murrelets (*Brachyramphus marmoratus*), with 2024 results supporting PAM as a cost-efficient alternative to traditional audio-visual surveys for assessing inland nesting habitat use in coastal forests. Beyond focal species, the program increasingly supports multi-taxa biodiversity monitoring through bioacoustic bycatch and soundscape analyses, linking species occurrence, habitat condition, and disturbance dynamics. Collectively, 2024 results demonstrate that PAM has reached operational maturity as the cornerstone of Phase II NWFP effectiveness monitoring. Sustaining adequate sampling effort and field capacity remains a challenge but essential, as reduced deployment intensity would increase uncertainty and weaken inference. Continued investment in PAM infrastructure and expertise is critical for informing adaptive management, supporting sustainable timber harvest, and evaluating ecosystem resilience under changing disturbance regimes.

## 1. Introduction

Northern spotted owl (*Strix occidentalis caurina*) population monitoring under the Northwest Forest Plan (NWFP) Interagency Effectiveness Monitoring Program continues to evolve from traditional field-based demography studies toward an integrated, range-wide framework based on a passive acoustic monitoring (PAM) network, remote landcover sensing and mapping, and machine learning. The PAM program provides a unified, statistically rigorous approach to assessing the status and trends of northern spotted owls, barred owls (*Strix varia*), and other forest wildlife across the NWFP area. This transition marks a critical inflection point in long-term monitoring, building upon more than three decades of spotted owl demographic research (Anthony et al. 2006, Forsman et al. 2011, Dugger et al. 2016, Franklin et al. 2021, Dugger et al. *in prep*). The current monitoring program uses new field survey techniques and analytical methods to produce landscape-scale inference on species occupancy, habitat associations, and trends in population size and distribution (Lesmeister and Jenkins 2022, Lesmeister et al. *in review*).

The PAM network was developed collaboratively by an interagency group of partners and now encompasses more than 4,000 survey stations where autonomous recording units (ARUs) are deployed annually across approximately 24 million acres of federally administered lands. These deployments follow a randomized 2+20% sample design (Lesmeister et al. 2021), generating data that are spatially representative of regional forest conditions and complementary to other NWFP monitoring modules (Davis et al. 2022a, Davis et al. 2022b). From 2018–2024, the network has accumulated nearly 8.5 million hours of acoustic recordings, analyzed with PNW-Cnet, a convolutional neural network model developed by the Pacific Northwest Bioacoustics Lab (Ruff et al. 2021, Ruff et al. 2023, Lesmeister et al. 2025b). These data provide the most extensive acoustic record of forest ecosystems in the Pacific Northwest. Regional survey designs often require tradeoffs be made between the number of sampling occasions and sites surveyed. The NWFP monitoring program has resolved many of these tradeoffs by using PAM with long-duration deployments, random site selection, multiscale clustered sampling, and high-throughput data processing for a wide range of species over large geographic regions.

Several new peer-reviewed and in-preparation analyses demonstrate that range-wide PAM has reached full operational maturity under the NWFP. Using data collected in 2023, Lesmeister et al. (*in review*) provided the first range-wide occupancy estimates for northern spotted owls and Jenkins et al. (2025) quantified complementary estimates for barred owls. Spotted owl detection probabilities exceeded 0.8 within four weeks of sampling when pairs were present and barred owl detection probabilities exceeded 0.9 after two weeks of sampling, confirming PAM as an efficient and scalable alternative to traditional call-back surveys associated with demographic monitoring. Critically, the findings collectively highlighted regional spotted owl extinction risks and potential mitigation measures that may reduce extinction likelihoods. The NWFP-wide predictions of barred owl distribution can directly inform the US Fish and Wildlife Service barred owl management strategy (USFWS 2024).

Building on these results, Lesmeister et al. (*in prep*) developed a six-step analytical framework to provide a structured, transparent approach for integrating territory-based and modern PAM-based occupancy estimates to support the methodological transition in the northern spotted owl monitoring program. This approach merges species occurrence data from these seemingly disparate datasets and modeling frameworks to produce integrated metrics of occupancy (Weldy et al. 2023). Initial findings indicate that persistent northern spotted owl use is concentrated in areas with low barred owl use and that coincide with fire refugia and mature forest structure, highlighting the value of integrating acoustic monitoring with spatial habitat indicators to inform conservation and management. These analyses collectively demonstrate that PAM can identify range-wide patterns of occupancy dynamics comparable in precision to historical demography studies, but at much broader spatial scales and provide information to guide conservation management decisions.

The effort summarized in this report extend and refine these findings, representing the next step in developing a comprehensive and integrated monitoring framework that links acoustic and habitat data to evaluate long-term ecosystem integrity and conservation effectiveness across the NWFP area. We extend PAM monitoring beyond northern spotted owls to include the proportion of monitoring hexagons with validated detections of barred owls and marbled murrelets (*Brachyramphus marmoratus*) for 2018–2024. We also report the estimated number of detections for all PNW-Cnet v5 sound classes by study area and state in 2024. Details on previous years’ results are available in earlier annual reports (Lesmeister et al. 2018, Lesmeister et al. 2019, Lesmeister et al. 2022, Lesmeister et al. 2023, Lesmeister et al. 2024).

## 2. Study Area

We collected PAM data within federally administered forest lands within the NWFP area (Fig. 1):

- 15 National Forests: Deschutes, Fremont-Winema, Gifford Pinchot, Klamath, Six Rivers, Mendocino, Mount Baker-Snoqualmie, Mount Hood, Okanogan-Wenatchee, Olympic, Rogue River-Siskiyou, Shasta-Trinity, Siuslaw, Umpqua, and Willamette
- 6 Bureau of Land Management Districts: Northern California, Medford, Lakeview, Coos Bay, Roseburg, and Northwest Oregon
- 8 National Park Lands: Golden Gate National Recreation Area, Muir Woods National Monument, Point Reyes National Seashore, Crater Lake National Park, Olympic National Park, North Cascades National Park, Mount Rainier National Park, and Redwood National and State Park
- Columbia River Gorge National Scenic Area that was administered by the US Forest Service

**Figure 1.**
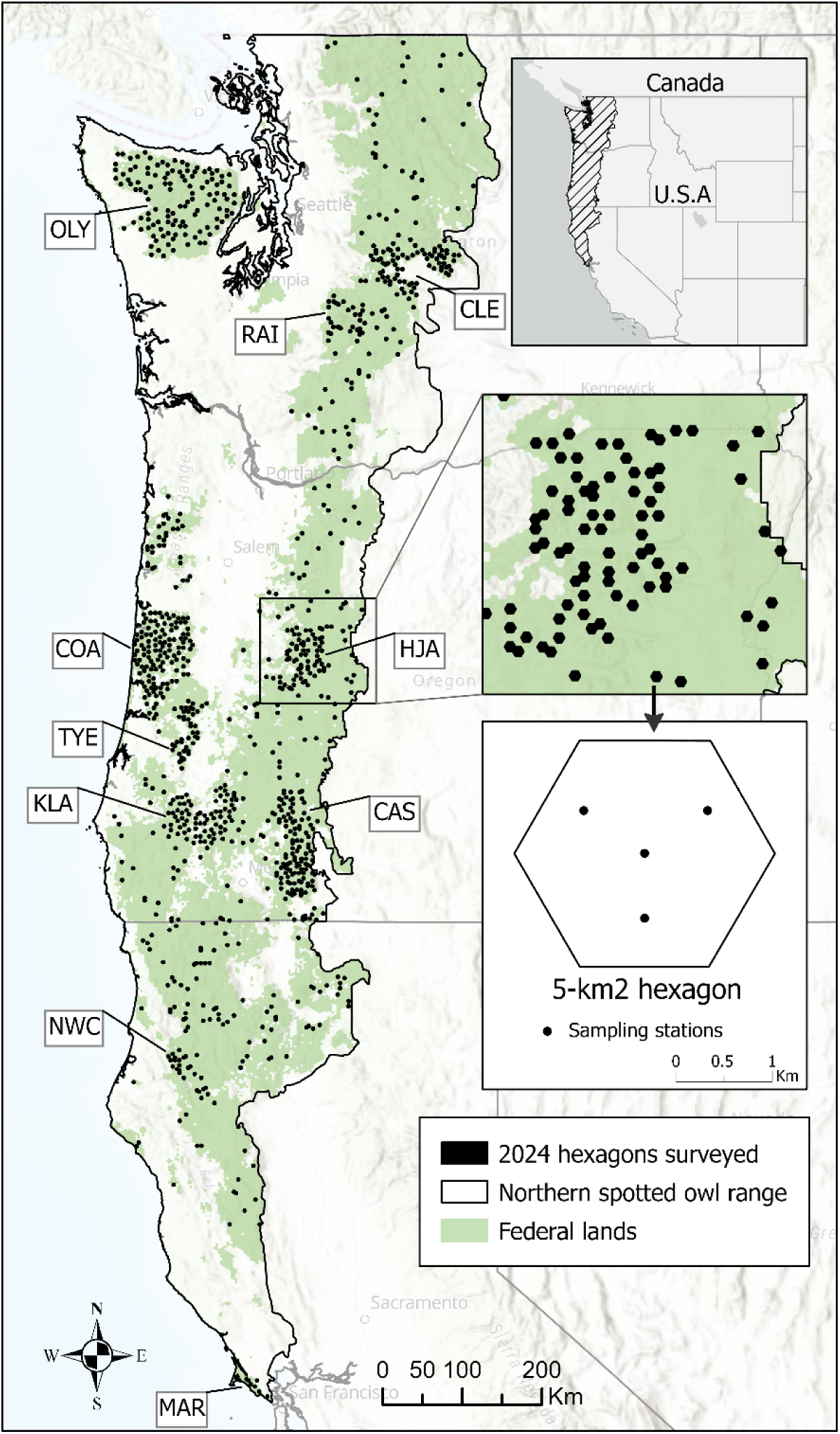
Locations of 5-km^2^ hexagons (*n* =1,020) surveyed on federal lands using passive acoustic monitoring in 2024. Inset shows typical placement of acoustic sampling stations. Historical study areas sampled at 20%: COA = Oregon Coast Range, OLY = Olympic Peninsula, KLA = Klamath Mountains, CLE = Cle Elum, TYE = Tyee, HJA = HJ Andrews Experimental Forest, CAS = Oregon South Cascades, NWC = Northwest California, MAR = Marin County, and RAI = Mount Rainier National Park. WA 2%, OR 2%, and CA 2% were data collected in each state on the 2% sampling outside the 20% sampling density on historical study areas.

Within the federal lands, we collected data within 10 historical northern spotted owl demographic study areas (Franklin et al. 2021). Nine of the study areas (OLY = Olympic Peninsula, WA; CLE = Cle Elum, WA; RAI = Mount Rainier National Park, WA; COA = Oregon Coast Range, OR; HJA = HJ Andrews Experimental Forest, OR; TYE = Tyee, OR; KLA = Klamath Mountains, OR; CAS = Oregon South Cascades, OR; NWC = Northwest California, CA) were long-term demographic study areas for northern spotted owl monitoring under the NWFP (Franklin et al. 2021). One study area (MAR = Marin County) was included due to long-term and ongoing northern spotted owl demographic monitoring (Fig. 1). In 2024, we deployed recording units within 48 designated wilderness areas (Table 1).

**Table 1.** The number of autonomous recording unit stations surveyed during 2024 in 48 designated wilderness areas administered by US Forest Service, US National Park Service, or US Bureau of Land Management.

| **Wilderness Area** | **Stations surveyed** |
| --- | --- |
| *US Forest Service* |  |
| Alpine Lakes | 27 |
| Boulder Creek | 1 |
| Buckhorn | 9 |
| Bull of the Woods | 3 |
| Clackamas | 2 |
| Clearwater | 2 |
| Colonel Bob | 8 |
| Cummins Creek | 3 |
| Devil's Staircase | 20 |
| Diamond Peak | 2 |
| Drift Creek | 2 |
| Glacier Peak | 12 |
| Goat Rocks | 8 |
| Indian Heaven | 2 |
| Kalmiopsis | 11 |
| Lake Chelan-Sawtooth | 4 |
| Marble Mountain | 8 |
| Mark O. Hatfield | 4 |
| Menagerie | 4 |
| Mount Hood | 9 |
| Mount Jefferson | 17 |
| Mount Skokomish | 4 |
| Mount Washington | 2 |
| Mountain Lakes | 10 |
| Noisy-Diobsud | 4 |
| North Fork Eel | 1 |
| Opal Creek | 10 |
| Pasayten | 12 |
| Red Buttes | 4 |
| Rock Creek | 1 |
| Rogue-Umpqua Divide | 10 |
| Salmon-Huckleberry | 1 |
| Siskiyou | 10 |
| Sky Lakes | 63 |
| Snow Mountain | 4 |
| The Brothers | 4 |
| Three Sisters | 49 |
| Trinity Alps | 19 |
| William O. Douglas | 23 |
| Yolla Bolly-Middle Eel | 4 |
| *US National Park Service* |  |
| Mount Rainier | 111 |
| Olympic | 238 |
| Phillip Burton | 7 |
| Stephen Mather | 22 |
| *US Bureau of Land Management* |  |
| Soda Mountain | 7 |
| Wild Rogue | 4 |
| Elkhorn Ridge | 4 |
| King Range | 3 |
| *TOTAL* | 789 |

## 6. Methods

### Sampling design

We created a uniform layer of 5-km^2^ hexagons that covered the entire range of the northern spotted owl (Lesmeister et al. 2021) that is available for download (USFWS 2021). This hexagon size is approximately the size of a northern spotted owl territory core area (Glenn et al. 2004, Schilling et al. 2013) and approximates the home range size reported for barred owls in the Pacific Northwest (Hamer et al. 2007, Singleton et al. 2010, Wiens et al. 2014). Within the historical demographic study areas, we randomly selected 20% of available hexagons that contained ≥50% forest capable lands and ≥25% federal ownership. Outside the historical demographic study areas, we randomly selected 2% of hexagons throughout the entire NWFP area following the same criteria for forest capable lands and federal ownership, stratified by physiographic province. Forest capable lands were those areas with suitable soil type, plant association, and elevation capable of developing into forest (Davis and Lint 2005). In a subset of our 20% study areas (OLY, COA, KLA), we surveyed non-adjacent hexagons to reduce the probability of detecting the same individual in multiple hexagons. Within each hexagon, we deployed 4 recorders (Fig 1). Combing the 2% and 20% randomized sample of 5-km^2^ hexagons created our 2+20% sampling design across the area of the Northwest Forest Plan.

We collected acoustic data at survey stations using Song Meter SM4s (*n* = 3,839) and Song Meter Mini (*n* = 213; Wildlife Acoustics, Maynard, MA), which are portable, weatherproof, and easily programmable ARUs. The SM4s had two built-in omni-directional microphones with signal-to-noise ratio of 80 decibels (dB) typical at 1 kilohertz (kHz), two SDHC/SDXC flash card slots, average of 543 h of recording battery life, and a recording bandwidth of 0.02–48.00 kHz at levels of -33.5–122.0 dB. We recorded with SM4s using only one microphone to reduce the amount of redundant data recorded and to extend recording time. The Song Meter Mini had the same bandwidth, signal-to-noise ratio of 78 dB, one omni-directional microphone, one SDHC/SDXC flash card slot, and 210–1040 h battery life depending on configuration. We programmed all recorders to use a sample rate of 32 kHz. These ARUs recorded sound with equivalent sensitivity to normal range of human hearing, and their effective listening radius may be affected by external factors such as terrain, vegetation, and weather events such as wind and rain. At each sampling station within a hexagon, we mounted ARUs to small trees (15–20 cm diameter at breast height) to allow microphones to extend past the bole for unobstructed recording ability. We deployed ARUs on federal land; mid-to-upper slope positions; ≥50 m from roads, trails, and streams to reduce vandalism and excessive noise; spaced ≥500 m apart; and located ≥200 m from edge of the hexagon. We programed ARUs to record from 1 h before sunset to 3 h after sunset, 2 h before sunrise to 2 h after sunrise, and for the first 10 min of every hour throughout the day and night.

### Data processing

Our multi-step workflow integrated PNW-Cnet v5 to efficiently process large volumes of audio data, combining automated identification and human validation (Ruff et al. 2021). PNW-Cnet v5 classifies call types for 84 species and had high precision for most sound classes (Lesmeister et al. 2025b). There were 108 sound classes with precision estimates >0.90 at the 0.95 threshold (recall ranging from 0.01–0.97). PNW-Cnet generates predictions—interpretable as confidence scores between 0.00–1.00—for each sound class for each 12 second clip (Ruff et al. 2020). We report the number of apparent detections, which are the number of clips with a score exceeding a classification score threshold (i.e., 0.95 for most target classes). We adjusted the number of apparent detections reported by model class precision (see Lesmeister et al. 2025b). We used a high classification threshold (0.95) to maximize precision over recall for this report.

We report the number of predicted detections for all PNW-Cnet v5 classes that had at least one predicted detection. Predicted detections for each sound class were calculated as the number of 12-second clips in the audio dataset to which the PNW-Cnet v5 assigned a score exceeding 0.95 for that class, multiplied by the precision estimate for that class.

### Data validation

Apparent detections of northern spotted owl, barred owl, and marbled murrelet call classes were validated by technicians using the program Kaleidoscope Pro to examine audio and associated spectrograms. We reviewed all clips with PNW-Cnet classification scores ≥0.50 for the northern spotted owl four-note location calls and ≥0.95 for survey tones (artificial pure tones played preceding or following call-back surveys) to maximize recall. We reviewed other northern spotted owl call type classes, the marbled murrelet keer call, and the barred owl eight-note territorial call at a classification threshold of ≥0.95, reviewing as many as necessary to validate weekly station-level occurrence. We did not review apparent detections for marbled murrelet keer calls in hexagons outside of the NWFP murrelet management zones (Raphael et al. 2018). Northern spotted owl detections within hexagons with ≥1 confirmed spotted owl call were reviewed by at least 2 technicians. During initial and secondary reviews, validators confirmed detection of species, identified sex using aspects of frequency and call duration (Dale et al. 2022), and identified artificial calls from call-back surveys where possible. We also reviewed apparent wolf (*Canus lupus*) detections with classification scores of ≥0.95.

Call-back surveys for northern spotted owls, barred owls, and, rarely, other species occurred in our study areas by biologists working on other research projects (e.g., Franklin et al. 2021, Wiens et al. 2021, Hobart et al. 2025) and project-level clearance surveys. These surveys broadcast recorded calls of northern spotted owls or other target species to elicit a territorial response. We distributed a recording consisting of a brief series of pure tones (1 s at 0.5, 1.5, and 1.0 kHz) for *Strix* call-back surveyors to voluntarily play at the same volume directly before or after northern spotted owl call-back surveys (USFWS 2021). We requested and received call-back survey information from most surveyors in or around our sampling locations after the recording season. We removed any validated detections of the surveyed species if there was a reported or suspected call-back survey in the hexagon on the same night.

We report the counts of validated detections of male and female northern spotted owl four-note location calls and survey tones and the proportion of surveyed hexagons with validated detections for northern spotted owls, marbled murrelets, and barred owls by study area and the 2% sample in each state.

### Background noise analysis

Background noise has consistently been found to be an important predictor of detection probabilities in species occurrence models (Duchac et al. 2020, Duchac et al. 2021, Rugg et al. 2025). Therefore, we used the sound pressure level analysis feature in Kaleidoscope Pro to quantify relative background noise levels at each sampling station which are available upon request but not reported here.

## 7 Results

The amount of data collected, and area surveyed within the NWFP area has increased each year of passive acoustic monitoring since 2018 (Table 2). In 2024, we surveyed a total of 4,052 stations in 1,020 hexagons (Table 2) with nearly 2.25 million h of recordings (Table 3) and approximately 1 petabyte of data collected. Deployments took place from February 26 – July 18, 2024. We collected data at 789 sampling stations in wilderness areas administered by US National Park Service (*n* = 378) or US Forest Service (*n* = 393) or US Bureau of Land Management (*n* = 18; Table 1). Each year we experience some degree of data loss by various sources such as wildfire, theft, animal damage, water intrusion, and software or hardware failure.

**Table 2.** Passive acoustic monitoring effort during 2018–2024 within in the Northwest Forest Plan Area, summarized by historical study areas (i.e., 20% sampling) and 2% sampling (outside 20% sample density) by state. Sampling in addition to 2% or 20% sampling indicated in parentheses. Historical study areas: COA = Oregon Coast Range, OLY = Olympic Peninsula, KLA = Klamath Mountains, CLE = Cle Elum, TYE = Tyee, HJA = HJ Andrews Experimental Forest, CAS = Oregon South Cascades, NWC = Northwest California, MAR = Marin County, and RAI = Mount Rainier National Park.

| Area | Number of hexagons |  |  |  |  |  |  | Number of stations |  |  |  |  |  |  |
| --- | --- | --- | --- | --- | --- | --- | --- | --- | --- | --- | --- | --- | --- | --- |
|  | 2018 | 2019 | 2020 | 2021 | 2022 | 2023 | 2024 | 2018<sup>a</sup> | 2019 | 2020 | 2021 | 2022 | 2023 | 2024 |
| COA | 120 | 106 | 120 | 120 | 117 | 120 | 120 | 578 | 413 | 471 | 475 | 466 | 476 | 479 |
| OLY | 88 | 120 | 119 | 119 | 120 | 120 | 118 | 436 | 472 | 464 | 472 | 474 | 479 | 472 |
| KLA |  | 63 | 73 | 73 | 73 | 73 | 73 |  | 244 | 290 | 276 | 292 | 292 | 291 |
| CLE |  |  | 69 | 75 | 75 | 75 | 75 |  |  | 267 | 298 | 299 | 300 | 300 |
| TYE |  |  |  | 40 | 40 | 40 | 40 |  |  |  | 155 | 160 | 160 | 160 |
| HJA |  |  |  | 70 | 70 | 70 | 69 |  |  |  | 279 | 280 | 278 | 273 |
| CAS |  |  |  | 98 | 98 | 98 | 98 |  |  |  | 380 | 391 | 391 | 389 |
| NWC |  |  |  | 30 | 30 | 30 | 30 |  |  |  | 120 | 119 | 118 | 117 |
| MAR |  |  |  | 7 | 7 | 7 | 7 |  |  |  | 26 | 27 | 27 | 28 |
| RAI |  |  |  | 11 | 24 | 32 | 28 |  |  |  | 43 | 96 | 128 | 111 |
| WA 2% |  |  |  |  | 23<sup>b</sup> | 90 | 81 |  |  |  |  | 91<sup>b</sup> | 355 | 323 |
| OR 2% |  |  |  |  | 13<sup>b</sup> | 178 | 186 |  |  |  |  | 52<sup>b</sup> | 709 | 732 |
| CA 2% |  |  |  |  |  | 76 | 95 |  |  |  |  |  | 299 | 377 |
| Total | 208 | 289 | 381 | 643 | 690 | 1,009 | 1,020 | 1,014 | 1,129 | 1,492 | 2,524 | 2,747 | 4,012 | 4,052 |

<sup>a</sup> During 2018 survey design was five stations per hexagon.
<sup>b</sup> In 2022, Gifford Pinchot National Forest in Washington and Umpqua National Forest in Oregon were sampled at 2%.

**Table 3.** Thousands of hours of passive acoustic monitoring data collected during 2018–2024 in each study area and processed for automated species identification with PNW-Cnet. Historical study areas: COA = Oregon Coast Range, OLY = Olympic Peninsula, KLA = Klamath Mountains, CLE = Cle Elum, TYE = Tyee, HJA = HJ Andrews Experimental Forest, CAS = Oregon South Cascades, NWC = Northwest California, MAR = Marin County, and RAI = Mount Rainier National Park. WA 2%, OR 2%, and CA 2% were data collected in each state on the 2% sampling outside the 20% sampling density on historical study areas.

| Area | 2018<sup>a</sup> | 2019 | 2020 | 2021 | 2022 | 2023 | 2024 |
| --- | --- | --- | --- | --- | --- | --- | --- |
| COA | 197 | 155 | 187 | 217 | 258 | 259 | 267 |
| OLY | 147 | 184 | 219 | 215 | 264 | 271 | 264 |
| KLA |  | 92 | 130 | 135 | 162 | 171 | 171 |
| CLE |  |  | 104 | 153 | 152 | 168 | 160 |
| TYE |  |  |  | 78 | 92 | 88 | 89 |
| HJA |  |  |  | 131 | 150 | 149 | 152 |
| CAS |  |  |  | 178 | 188 | 195 | 199 |
| NWC |  |  |  | 48 | 54 | 62 | 62 |
| MAR |  |  |  | 11 | 10 | 15 | 15 |
| RAI |  |  |  | 11 | 51 | 71 | 56 |
| WA 2% |  |  |  |  | 44<sup>b</sup> | 192 | 186 |
| OR 2% |  |  |  |  | 30<sup>b</sup> | 385 | 420 |
| CA 2% |  |  |  |  |  | 151 | 204 |
| Total | 344 | 431 | 640 | 1,177 | 1,455 | 2,177 | 2,245 |

<sup>a</sup> During 2018 survey design was five stations per hexagon.
<sup>b</sup> In 2022, Gifford Pinchot National Forest in Washington and Umpqua National Forest in Oregon were sampled at 2%.

In 2024, we received broadcast survey occurrence records for barred owl, northern spotted owl, American goshawk (*Astur atricapillus*), sharp-shinned hawk (*Accipiter striatus*), northern saw-whet owl (*Aegolius acadicus*), great horned owl (*Bubo virginianus*), and osprey (*Pandion haliaetus*) call-back surveys within or adjacent to our sampled hexagons. Northern spotted owl broadcast surveys were reported in 19% of our hexagons (*n* = 197). We removed northern spotted owl calls in 170 hexagons due to overlapping broadcast records. There were 40 hexagons (4%) with reported barred owl broadcast surveys (Oregon and California), and we removed detections overlapping surveys from 5 hexagons. The NWC and KLA study areas had the highest proportion of hexagons with broadcast surveys reported: 0.73 (41 of 73 hexagons) and 0.56 (22 of 30 hexagons), respectively. No call-back surveys were reported in CLE, MAR, or RAI. We confirmed artificial survey tones in 116 hexagons across all states (11%), but found none in CLE, MAR, OLY, or RAI (Table 4).

**Table 4.** Predicted number of detections of PNW-Cnet v5 sound classes from passive acoustic monitoring data by study area collected in 2024. Predicted detections for each sound class were calculated as the number of 12-second clips in the audio dataset to which the PNW-Cnet v5 assigned a score exceeding 0.95 for that class, multiplied by the precision estimate. Validated call counts are given for the primary northern spotted owl call (NSO; after call-back survey removal) and survey tone call classes which were fully reviewed at 0.5 and 0.95 thresholds, respectively. We also validated wolf detections. Classes with zero apparent detections not shown. Historical study areas (20% sampling): CAS = Southwest Cascades, OR; CLE = Cle Elum, WA; COA = Coast Range, OR; HJA = HJ Andrews Experimental Forest, OR; KLA = Klamath Mountains, OR and CA; MAR = Marin County, CA; NWC = Northwest California, CA; OLY = Olympic Peninsula, WA; RAI = Mount Rainier National Park, WA; and TYE = Tyee, OR. 2% random sample of federal lands in Washington (WA 2%), Oregon (OR 2%), and California (CA 2%).

| Sound | CAS | CLE | COA | HJA | KLA | MAR | NWC | OLY | RAI | TYE | CA 2% | OR 2% | WA 2% |
| --- | --- | --- | --- | --- | --- | --- | --- | --- | --- | --- | --- | --- | --- |
| NSO female 4-note<sup>a</sup> | 42 | 10 | 59 | 70 | 35 | 482<sup>b</sup> | 118 | 39 | 0 | 0 | 380 | 133 | 6 |
| NSO male 4-note<sup>a</sup> | 1,654 | 883 | 60 | 1,274 | 1,005 | 6623<sup>b</sup> | 4,046 | 2,872 | 26 | 10 | 1,275 | 3,255 | 314 |
| NSO series | 43 | 9 | 28 | 24 | 15 | 1,374 | 131 | 40 | 0 | 1 | 2,149 | 320 | 7 |
| NSO bark | 93 | 3 | 2 | 24 | 53 | 254 | 137 | 81 | 0 | 1 | 533 | 55 | 0 |
| *Strix* owl contact whistle | 1,285 | 110 | 926 | 337 | 2,526 | 766 | 2,378 | 433 | 104 | 707 | 762 | 3,147 | 284 |
| Survey tone<sup>a</sup> | 36 | 0 | 119 | 3 | 257 | 0 | 19 | 0 | 0 | 49 | 18 | 48 | 8 |
| Barred owl 8-note | 10,567 | 7,175 | 59,905 | 13,368 | 20,129 | 47 | 5,636 | 26,371 | 5,409 | 15,458 | 2,949 | 46,654 | 15,018 |
| Barred owl inspection | 10,146 | 10,195 | 35,236 | 7,883 | 23,801 | 312 | 1,869 | 9,131 | 3,090 | 12,141 | 2,546 | 30,633 | 13,967 |
| Barred owl series | 1,163 | 425 | 9,544 | 1,383 | 3,226 | 3 | 269 | 2,260 | 319 | 3,275 | 422 | 6,018 | 1,421 |
| Marbled murrelet keer | 49 | 72 | 44,847 | 131 | 172 | 29 | 31 | 34,915 | 361 | 11 | 1,935 | 4,831 | 3,281 |
| Airplane | 1 | 41 | 93 | 4 | 44 | 5 | 0 | 1 | 2 | 7 | 128 | 1,342 | 136 |
| Chainsaw | 303 | 12,021 | 7,416 | 1,369 | 6,186 | 23 | 2,444 | 2,005 | 1 | 2,287 | 2,214 | 5,956 | 1,303 |
| Military jet | 1,038 | 20 | 1,991 | 121 | 1,333 | 0 | 3,275 | 14 | 33 | 0 | 32,497 | 769 | 118 |
| Gunshot | 520 | 1,765 | 2,318 | 136 | 876 | 68 | 127 | 2,888 | 183 | 442 | 570 | 8,017 | 3,654 |
| Highway noise | 1,032 | 3,987 | 60 | 192 | 8,478 | 0 | 30 | 1,110 | 2 | 18 | 200 | 1,301 | 122 |
| Human speech | 1,157 | 794 | 2,302 | 725 | 922 | 1,680 | 417 | 1,041 | 595 | 467 | 1,567 | 5,176 | 928 |
| Train | 0 | 525 | 626 | 1 | 2,730 | 2 | 0 | 1 | 0 | 2 | 2,236 | 900 | 573 |
| Yarder | 133 | 13 | 64,679 | 8,213 | 36,006 | 47 | 6 | 12,989 | 0 | 54,764 | 1,044 | 58,774 | 341 |
| American crow | 81 | 114 | 356 | 86 | 442 | 995 | 40 | 140 | 41 | 426 | 283 | 571 | 10 |
| Common raven | 89,001 | 81,989 | 155,316 | 24,086 | 144,838 | 40,129 | 24,151 | 47,443 | 9,199 | 76,111 | 140,318 | 222,316 | 47,058 |
| Steller's jay | 319,263 | 38,319 | 551,948 | 245,026 | 957,007 | 31,657 | 287,996 | 113,577 | 15,729 | 252,627 | 678,311 | 1,018,858 | 46,014 |
| Clark's nutcracker | 2,285 | 2,988 | 5 | 24 | 16 | 1 | 8 | 5 | 283 | 8 | 1,172 | 8,670 | 7,444 |
| Canada jay | 7,408 | 3,523 | 16,213 | 6,289 | 3,819 | 7 | 141 | 17,598 | 6,670 | 5,575 | 1,179 | 18,276 | 16,359 |
| Bullfrog | 20 | 1 | 4,326 | 0 | 2,208 | 11 | 0 | 0 | 0 | 2,421 | 705 | 10,525 | 1 |
| Chicken | 96 | 253 | 3,613 | 6 | 26,362 | 1,889 | 255 | 811 | 13 | 2,215 | 3,567 | 11,255 | 34 |
| Cow | 10 | 0 | 17 | 0 | 17 | 4 | 0 | 0 | 0 | 15 | 15 | 31 | 1 |
| Stream noise | 158 | 789 | 45,765 | 14 | 102 | 2,238 | 8 | 20 | 1 | 6,512 | 289 | 2,389 | 5,937 |
| Cricket | 14,079 | 28,360 | 24,507 | 126,904 | 1,613,511 | 2,486 | 312,316 | 5,283 | 530 | 214,548 | 1,813,146 | 1,319,833 | 42,716 |
| Dog | 512 | 5,299 | 4,560 | 118 | 52,021 | 6,020 | 3,486 | 976 | 1 | 6,979 | 8,230 | 25,318 | 1,073 |
| Buzzing insect | 810,371 | 583,895 | 224,066 | 286,870 | 499,014 | 3,776 | 169,038 | 518,703 | 156,357 | 131,598 | 633,358 | 1,121,359 | 692,763 |
| Frog chorus | 689,558 | 289,046 | 309,918 | 184,966 | 249,702 | 51,502 | 117,794 | 280,986 | 48,655 | 329,940 | 540,901 | 819,246 | 144,917 |
| Rain | 7 | 251 | 4,501 | 1,526 | 36 | 0 | 1,217 | 8,653 | 625 | 989 | 2,697 | 4,564 | 2,020 |
| Thunder | 0 | 0 | 0 | 0 | 5 | 0 | 0 | 122 | 0 | 0 | 0 | 9 | 0 |
| Tree creaking | 11,297 | 40,573 | 13,429 | 1,931 | 6,176 | 154 | 633 | 3,548 | 2,953 | 7,422 | 7,682 | 18,201 | 12,052 |
| Coyote | 0 | 1 | 3 | 0 | 4 | 3 | 0 | 1 | 0 | 0 | 2 | 2 | 0 |
| Wolf howl<sup>a</sup> | 0 | 1 | 0 | 0 | 0 | 0 | 0 | 0 | 0 | 0 | 0 | 0 | 0 |
| American pika | 7,579 | 10,094 | 14 | 3,475 | 41 | 0 | 14 | 17 | 2,874 | 6 | 465 | 1,902 | 17,733 |
| Deer snort | 14 | 4 | 22 | 16 | 48 | 1 | 19 | 25 | 2 | 10 | 22 | 56 | 4 |
| Douglas squirrel rattle | 46,532 | 61,209 | 54,839 | 15,070 | 34,064 | 140 | 31,434 | 55,273 | 12,740 | 17,289 | 75,311 | 74,788 | 70,120 |
| **Douglas squirrel chirp** | 43,425 | 38,871 | 26,427 | 13,459 | 13,554 | 7 | 22,593 | 44,915 | 13,052 | 9,638 | 57,252 | 55,610 | 38,378 |
| **Chipmunk spp. chirp** | 89,905 | 105,081 | 143,077 | 59,095 | 140,416 | 675 | 30,223 | 34,144 | 20,152 | 37,673 | 127,824 | 238,185 | 76,158 |
| **Black bear juvenile** | 7 | 9 | 14 | 1 | 9 | 0 | 3 | 1 | 2 | 2 | 13 | 83 | 22 |
| **Sandhill crane** | 86 | 0 | 0 | 0 | 0 | 0 | 0 | 0 | 0 | 0 | 0 | 33 | 0 |
| **Canada goose** | 20,706 | 2,825 | 7,298 | 1,299 | 2,150 | 163 | 70 | 555 | 270 | 1,704 | 798 | 15,543 | 459 |
| **California quail** | 0 | 0 | 0 | 0 | 18 | 540 | 0 | 0 | 0 | 1 | 9 | 4 | 0 |
| **Common nighthawk "peent"** | 510,897 | 73,580 | 66,980 | 59,418 | 152,814 | 18 | 8,003 | 123,057 | 3,741 | 61,060 | 182,936 | 931,700 | 239,162 |
| **Common nighthawk "boom"** | 351,823 | 20,102 | 43,929 | 22,995 | 129,546 | 23 | 4,692 | 30,003 | 13 | 39,654 | 123,613 | 582,558 | 107,655 |
| **Sooty grouse hoot** | 85,713 | 114,887 | 22,334 | 271,676 | 46,575 | 4 | 3,015 | 964,279 | 21,385 | 803,328 | 21,712 | 370,430 | 134,406 |
| **Sooty grouse call** | 6 | 3 | 22 | 16 | 27 | 0 | 10 | 28 | 0 | 9 | 3 | 14 | 28 |
| **Wilson's snipe** | 12 | 4 | 0 | 0 | 0 | 0 | 0 | 1 | 0 | 0 | 1 | 3 | 0 |
| **Wild turkey** | 551 | 33 | 11 | 19 | 2,038 | 393 | 0 | 5 | 0 | 99 | 1,172 | 1,148 | 42 |
| **Mountain quail song** | 22,748 | 21 | 30,614 | 8,251 | 101,943 | 1 | 108,783 | 10 | 4 | 44,783 | 260,275 | 167,201 | 25 |
| **Mountain quail call** | 330 | 10 | 82 | 69 | 390 | 0 | 393 | 11 | 0 | 156 | 3,531 | 590 | 160 |
| **Band-tailed pigeon song** | 7,052 | 876 | 614,219 | 13,036 | 58,682 | 37,593 | 6,287 | 74,168 | 1,168 | 41,484 | 44,110 | 155,601 | 4,091 |
| **Band-tailed pigeon grunt** | 3 | 4 | 69 | 10 | 40 | 7 | 3 | 140 | 8 | 6 | 30 | 120 | 13 |
| **Common poorwill** | 98,765 | 199,174 | 19 | 206 | 13,463 | 0 | 21,980 | 7 | 1 | 31 | 112,118 | 310,657 | 106,833 |
| **Eurasian collared dove** | 0 | 0 | 0 | 0 | 35 | 0 | 0 | 0 | 0 | 0 | 0 | 6 | 0 |
| **Mourning dove** | 1,414 | 495 | 913 | 360 | 22,871 | 11,242 | 24,078 | 86 | 0 | 2,583 | 76,721 | 99,010 | 2,151 |
| **Northern saw-whet owl** | 154,140 | 28,273 | 353,412 | 47,473 | 31,410 | 21,648 | 45,461 | 78,041 | 6,849 | 206,119 | 90,418 | 209,513 | 14,828 |
| **Long-eared owl** | 8 | 4 | 0 | 0 | 0 | 0 | 1 | 0 | 0 | 0 | 1 | 7 | 1 |
| **Great horned owl song** | 45,756 | 86,683 | 4,797 | 2,203 | 40,432 | 8,403 | 3,002 | 4,290 | 110 | 14,140 | 42,353 | 53,432 | 30,451 |
| **Great horned owl calls** | 499 | 1,585 | 414 | 23 | 293 | 50 | 254 | 101 | 1 | 125 | 2,752 | 2,091 | 208 |
| Northern<br>pygmy-owl | 35,547 | 37,757 | 988,484 | 118,979 | 85,269 | 258 | 32,632 | 111,627 | 11,468 | 430,192 | 118,947 | 449,095 | 40,272 |
| Western<br>screech-owl<br>trill | 6,899 | 948 | 70,331 | 4,279 | 285,127 | 89 | 28,057 | 8,818 | 1 | 108,125 | 73,323 | 132,032 | 1,579 |
| Western<br>screech-owl<br>bark | 3 | 2 | 1 | 1 | 37 | 0 | 13 | 2 | 0 | 3 | 21 | 12 | 1 |
| Western<br>screech-owl<br>juvenile | 142 | 7,911 | 2,680 | 3,084 | 123 | 22 | 1,385 | 1,409 | 106 | 3,277 | 1,516 | 7,247 | 5,991 |
| Flammulated<br>owl | 5,409 | 160,137 | 9 | 3 | 177 | 0 | 39,054 | 8 | 66 | 154 | 80,731 | 112,040 | 24,914 |
| Great gray owl<br>song | 1,022 | 60 | 313 | 1,178 | 1,193 | 1 | 5 | 13 | 1 | 26 | 214 | 696 | 36 |
| Great gray owl<br>calls | 0 | 0 | 0 | 0 | 0 | 0 | 0 | 0 | 0 | 0 | 0 | 18 | 0 |
| Cooper's hawk<br>series call | 35 | 0 | 5 | 0 | 2 | 0 | 2 | 0 | 0 | 7 | 1 | 2 | 0 |
| American<br>goshawk series | 4,430 | 1,070 | 73 | 213 | 364 | 21 | 2,961 | 262 | 57 | 54 | 1,343 | 1,420 | 1,223 |
| American<br>goshawk<br>scream | 3,564 | 2,182 | 432 | 1,191 | 4,129 | 267 | 763 | 1,347 | 757 | 59 | 6,099 | 19,098 | 14,250 |
| Sharp-shinned<br>hawk series<br>call | 1 | 0 | 1 | 0 | 1 | 0 | 0 | 0 | 1 | 0 | 1 | 0 | 5 |
| Merlin | 8 | 4 | 38 | 3 | 0 | 1 | 1 | 538 | 0 | 1 | 0 | 8 | 6 |
| American<br>kestrel | 134 | 77 | 130 | 36 | 152 | 1 | 19 | 313 | 54 | 151 | 219 | 520 | 459 |
| Raptor spp.<br>scream call | 1 | 1 | 93 | 1 | 51 | 0 | 0 | 4 | 60 | 30 | 53 | 16 | 1 |
| Hermit thrush<br>song | 1,947,284 | 1,392,190 | 52,257 | 834,909 | 2,091,214 | 31,517 | 218,936 | 212,379 | 315,347 | 452,350 | 699,820 | 2,192,774 | 1,494,968 |
| Hermit thrush<br>chit-chit call | 1 | 0 | 0 | 0 | 2 | 0 | 0 | 0 | 0 | 0 | 0 | 3 | 3 |
| Hermit thrush<br>wheeze | 32,889 | 9,575 | 1,087 | 9,115 | 13,119 | 1,371 | 1,165 | 2,162 | 2,300 | 6,048 | 7,750 | 15,266 | 8,215 |
| Swainson's<br>thrush song | 97,607 | 142,845 | 7,473,379 | 902,153 | 300,512 | 161,188 | 136 | 253,762 | 14,189 | 217,625 | 39,235 | 2,558,465 | 488,319 |
| Swainson's<br>thrush call | 89 | 211 | 122,335 | 2,873 | 1,076 | 2,083 | 19 | 1,878 | 71 | 1,377 | 153 | 23,147 | 1,025 |
| Olive-sided<br>flycatcher song | 166,113 | 135,105 | 50,537 | 88,637 | 106,434 | 32,290 | 64,264 | 137,229 | 7,678 | 18,640 | 258,167 | 388,491 | 165,581 |
| Olive-sided<br>flycatcher call | 41,193 | 23,239 | 9,749 | 13,996 | 34,334 | 1,486 | 4,584 | 46,798 | 10,299 | 7,939 | 57,461 | 76,679 | 42,442 |
| Wrentit | 3,608 | 1 | 133,606 | 14 | 195,310 | 47,626 | 3,070 | 13 | 0 | 28,611 | 46,560 | 49,247 | 0 |
| Western wood-pewee | 2,397 | 9,563 | 2,555 | 839 | 4,326 | 36 | 2,462 | 2,362 | 56 | 1,791 | 14,468 | 16,345 | 15,696 |
| Dusky flycatcher | 11,190 | 4,994 | 26 | 21 | 115 | 0 | 2,974 | 12 | 3 | 24 | 13,615 | 16,711 | 1,971 |
| Purple finch | 419 | 998 | 165 | 15 | 85 | 85 | 49 | 63 | 0 | 127 | 294 | 351 | 156 |
| Varied thrush song | 16,451 | 299,505 | 791,590 | 229,979 | 7,045 | 17 | 796 | 2,093,716 | 352,578 | 97,679 | 10,498 | 437,147 | 959,240 |
| Varied thrush call | 1,223 | 776 | 1,225 | 697 | 1,475 | 187 | 846 | 3,615 | 957 | 541 | 4,582 | 3,042 | 2,404 |
| Dark-eyed junco | 7,997 | 12,003 | 276 | 4,907 | 2,733 | 62 | 2,906 | 15,693 | 1,548 | 977 | 4,482 | 8,658 | 9,342 |
| Red crossbill | 0 | 0 | 64 | 0 | 5 | 0 | 0 | 11 | 0 | 0 | 1 | 47 | 22 |
| Townsend's solitaire | 93,055 | 39,667 | 1,314 | 42,624 | 44,977 | 43 | 16,038 | 12,774 | 1,953 | 2,692 | 86,541 | 121,160 | 37,903 |
| Western tanager call | 0 | 0 | 0 | 0 | 1 | 0 | 2 | 0 | 0 | 0 | 0 | 7 | 0 |
| Spotted towhee call | 6,975 | 234 | 5,977 | 1,044 | 18,354 | 9,050 | 22,787 | 24 | 1 | 8,051 | 59,949 | 29,166 | 964 |
| Chickadee song | 189,692 | 13,864 | 12 | 201 | 203 | 0 | 6,005 | 9 | 45 | 29 | 48,251 | 184,184 | 8,644 |
| Chickadee spp. call | 3,097 | 1,163 | 0 | 1 | 0 | 0 | 149 | 0 | 21 | 0 | 1,736 | 3,891 | 767 |
| Mountain bluebird | 87 | 1,556 | 0 | 1 | 0 | 0 | 1 | 1 | 0 | 0 | 1 | 65 | 50 |
| Nuthatch spp. song | 3,390,537 | 1,024,689 | 363,319 | 880,186 | 2,047,695 | 20,184 | 361,937 | 828,567 | 214,694 | 769,200 | 1,518,938 | 3,264,897 | 1,105,163 |
| Nuthatch spp. call | 954 | 364 | 20 | 445 | 249 | 0 | 128 | 127 | 95 | 110 | 220 | 703 | 457 |
| American robin whinny | 10,492 | 5,516 | 14,575 | 8,391 | 8,389 | 509 | 2,046 | 22,456 | 457 | 10,995 | 5,483 | 22,440 | 4,403 |
| American robin other calls | 3,346 | 880 | 3,856 | 2,205 | 3,383 | 44 | 1,937 | 4,709 | 98 | 2,386 | 3,210 | 6,870 | 1,394 |
| Hutton's vireo | 0 | 0 | 15 | 0 | 4 | 0 | 3 | 22 | 0 | 3 | 10 | 14 | 0 |
| Northern flicker series call | 63,484 | 30,099 | 29,307 | 28,481 | 42,212 | 2,524 | 15,329 | 10,146 | 1,284 | 73,992 | 53,425 | 93,641 | 11,662 |
| Northern flicker "skew" | 14,229 | 5,150 | 4,262 | 4,236 | 21,301 | 210 | 4,787 | 6,580 | 772 | 8,733 | 16,072 | 22,534 | 6,495 |
| Downy woodpecker | 3 | 5 | 2 | 0 | 55 | 1 | 4 | 10 | 0 | 46 | 135 | 42 | 38 |
| Woodpecker drum (non-sapsucker) | 21,195 | 12,983 | 9,516 | 5,155 | 6,990 | 1,135 | 3,650 | 11,488 | 668 | 9,819 | 13,349 | 29,088 | 5,590 |
| Pileated woodpecker | 26,379 | 7,980 | 45,381 | 11,474 | 32,575 | 3,781 | 6,974 | 12,836 | 443 | 23,617 | 28,447 | 61,149 | 5,947 |
| White-headed woodpecker | 37 | 8 | 5 | 0 | 1 | 0 | 77 | 0 | 0 | 0 | 218 | 41 | 35 |
| Hairy woodpecker | 339 | 121 | 342 | 318 | 262 | 43 | 125 | 234 | 41 | 146 | 633 | 689 | 223 |
| Acorn woodpecker | 11 | 2 | 3 | 2 | 3,026 | 22 | 2,964 | 1 | 25 | 0 | 2,786 | 1,466 | 1 |
| Sapsucker spp. drum | 4,913 | 1,172 | 6,937 | 4,815 | 825 | 6 | 705 | 1,186 | 2,106 | 3,852 | 1,346 | 4,634 | 1,065 |

<sup>a</sup> Validated counts from full review at the 0.5 threshold or at a 0.95 threshold for Survey\_Tone and wolf howl.
<sup>b</sup> To balance effort amidst high calling rates, caller sex was not verified within every clip. Instead, we confirmed $\geq 1$ occurrence of both male and female calling weekly at each recording station.

### Focal species detections

In 2024 we observed a wide range number of detections of northern spotted owl in 20% and 2% sample areas. As in all previous years, we found more male spotted owls than female spotted owl detections except for COA (Table 4). California study areas (MAR, NWC) had the highest proportion of hexagons with spotted owl detections (Table 5). Study areas in the Washington Cascades (CLE, RAI, WA 2%) and two study areas in Oregon (TYE and COA) had some of the lowest proportion of hexagons with detections (Table 5). The proportion of hexagons with detections fluctuated in most study areas with multiple years of surveys, but we observed large declines in the hexagons with detections in TYE and KLA study areas (Table 5). The TYE study area has declined from 0.38 (15 hexagons) to 0.05 (2 hexagons) from 2021–2024. While KLA has declined by roughly half since 2022 from 51% (37 hexagons) to 23% (17 hexagons) 2024 (Table 5). We did not detect northern spotted owls in the Columbia River Gorge National Scenic Area, Deschutes National Forest, or North Cascades National Park. We also did not detect any northern spotted owl north of Stevens Pass in the Washington Cascades (30 hexagons).

**Table 5.** Proportion of monitored hexagons with validated detections of northern spotted owls for years that surveys were conducted (2018–2024) within the Northwest Forest Plan Area. Historical study areas sampled at 20%: COA = Oregon Coast Range, OLY = Olympic Peninsula, KLA = Klamath Mountains, CLE = Cle Elum, TYE = Tyee, HJA = HJ Andrews Experimental Forest, CAS = Oregon South Cascades, NWC = Northwest California, MAR = Marin County, and RAI = Mount Rainier National Park. WA 2%, OR 2%, and CA 2% were data collected in each state on the 2% sampling outside the 20% sampling density on historical study areas.

| Area | 2018 | 2019 | 2020 | 2021 | 2022 | 2023 | 2024 |
| --- | --- | --- | --- | --- | --- | --- | --- |
| COA | 0.16 | 0.13 | 0.09 | 0.15 | 0.09 | 0.11 | 0.10 |
| OLY | 0.16 | 0.17 | 0.24 | 0.19 | 0.27 | 0.25 | 0.14 |
| KLA |  | 0.43 | 0.53 | 0.49 | 0.51 | 0.37 | 0.23 |
| CLE |  |  | 0.19 | 0.19 | 0.11 | 0.20 | 0.13 |
| TYE |  |  |  | 0.38 | 0.28 | 0.15 | 0.05 |
| HJA |  |  |  | 0.39 | 0.39 | 0.43 | 0.35 |
| CAS |  |  |  | 0.35 | 0.31 | 0.23 | 0.31 |
| NWC |  |  |  | 0.77 | 0.73 | 0.80 | 0.60 |
| MAR |  |  |  | 1.00 | 1.00 | 1.00 | 1.00 |
| RAI |  |  |  | 0.09 | 0.04 | 0.06 | 0.07 |
| WA 2% |  |  |  |  | 0.00<sup>a</sup> | 0.09 | 0.07 |
| OR 2% |  |  |  |  | 0.54<sup>a</sup> | 0.24 | 0.17 |
| CA 2% |  |  |  |  |  | 0.37 | 0.38 |

<sup>a</sup> In 2022, Gifford Pinchot National Forest in Washington and Umpqua National Forest in Oregon were sampled at 2%.

We detected barred owl calling in 84% of surveyed hexagons (Table 6; Figure 2). In Washington, we detected barred owls in 90% of surveyed hexagons (*n* = 321) with a slightly lower rate (73%) in CLE (Table 6). In Oregon, we detected barred owls in 92% of hexagons (*n* = 561; Figure 2). The greatest numbers of barred owl 8-note calls were recorded in COA and the Oregon 2% areas (Table 4). We found barred owls in 50% of California hexagons (*n* = 78) and CA study areas had lower proportion of hexagons and numbers of barred owl 8-note calls compared to sites in Oregon and Washington (Table 4, Table 6). Mendocino National Forest in California was the only surveyed federal management unit with no barred owl calls detected.

**Figure 2.**
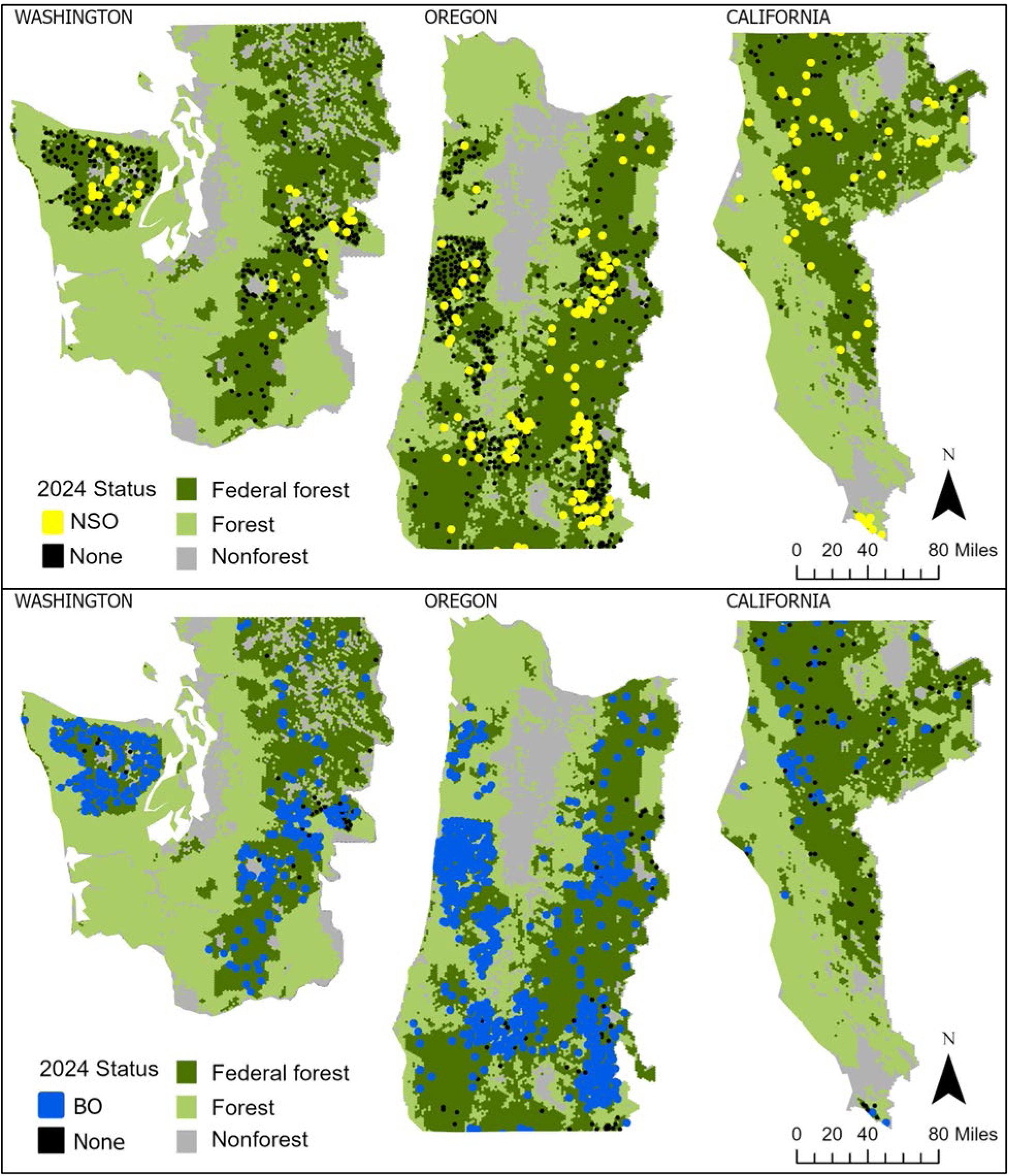
Locations of 5-km^2^ hexagons with validated northern spotted owl (NSO: *n* = 213) and barred owl (BO; *n* = 857) detections in 2024 acoustic monitoring.

**Table 6.** Proportion of monitoring hexagons with validated detections of barred owls for years that surveys were conducted (2018–2024) within the Northwest Forest Plan Area. Historical study areas sampled at 20%: COA = Oregon Coast Range, OLY = Olympic Peninsula, KLA = Klamath Mountains, CLE = Cle Elum, TYE = Tyee, HJA = HJ Andrews Experimental Forest, CAS = Oregon South Cascades, NWC = Northwest California, MAR = Marin County, and RAI = Mount Rainier National Park. WA 2%, OR 2%, and CA 2% were data collected in each state on the 2% sampling outside the 20% sampling density on historical study areas.

| Area | 2018 | 2019 | 2020 | 2021 | 2022<sup>a</sup> | 2023 | 2024 |
| --- | --- | --- | --- | --- | --- | --- | --- |
| COA | 0.99 | 0.99 | 1.00 | 0.99 | 0.75 | 0.99 | 1.00 |
| OLY | 0.92 | 0.93 | 0.93 | 0.90 | 0.60 | 0.93 | 0.95 |
| KLA |  | 0.86 | 0.95 | 0.95 | 0.73 | 0.97 | 0.92 |
| CLE |  |  | 0.77 | 0.65 | 0.35 | 0.80 | 0.73 |
| TYE |  |  |  | 1.00 | 0.85 | 1.00 | 1.00 |
| HJA |  |  |  | 0.99 | 0.57 | 0.97 | 0.96 |
| CAS |  |  |  | 0.95 | 0.43 | 0.94 | 0.93 |
| NWC |  |  |  | 0.73 | 0.33 | 0.67 | 0.67 |
| MAR |  |  |  | 0.29 | 0.14 | 0.14 | 0.29 |
| RAI |  |  |  | 0.82 | 0.46 | 0.94 | 0.96 |
| WA 2% |  |  |  |  | 0.65<sup>b</sup> | 0.93 | 0.89 |
| OR 2% |  |  |  |  | 0.46<sup>b</sup> | 0.84 | 0.83 |
| CA 2% |  |  |  |  |  | 0.37 | 0.33 |

<sup>a</sup> In 2022, we didn't extensively validate barred owl classes, thus adjusted PNW-Cnet model predicted proportions are reported.
<sup>b</sup> In 2022, only Gifford Pinchot National Forest in Washington and Umpqua National Forest in Oregon were sampled at 2%.

Marbled murrelet keer calls were identified in 52% of hexagons within NWFP marbled murrelet management zones (Figure 3). Marbled murrelets were commonly detected (87-97% of hexagons) in the two most coastal 20% study areas (COA, OLY; Table 7; Figure 3). A higher proportion of the OR 2% sampled areas had murrelet detections compared to WA 2% and CA 2%, where sampling was further from coastal forest (Figure 3; Table 7).

**Figure 3.**
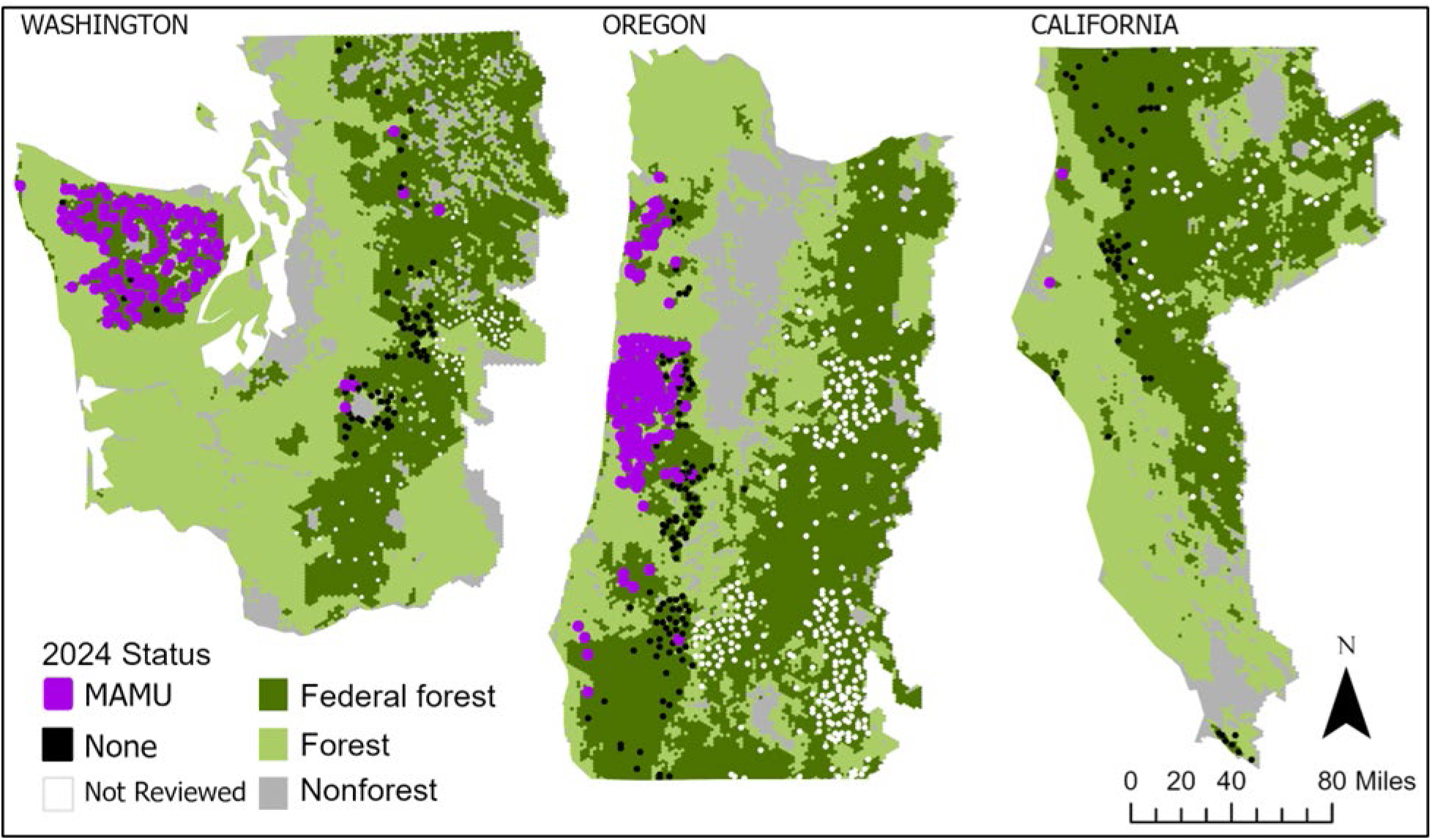
Locations of 5-km^2^ hexagons with validated marbled murrelet (MAMU; *n* = 268) detections in 2024. Only hexagons within the marbled murrelet Northwest Forest Plan management zones were reviewed (*n* = 511).

**Table 7.** Proportion of monitoring hexagons with validated detections of marbled murrelets for years that surveys were conducted (2018–2024) within the Northwest Forest Plan Area. Historical study areas sampled at 20%: COA = Oregon Coast Range, OLY = Olympic Peninsula, KLA = Klamath Mountains, CLE = Cle Elum, TYE = Tyee, HJA = HJ Andrews Experimental Forest, CAS = Oregon South Cascades, NWC = Northwest California, MAR = Marin County, and RAI = Mount Rainier National Park. WA 2%, OR 2%, and CA 2% were data collected in each state on the 2% sampling outside the 20% sampling density on historical study areas.

| Area | 2018 | 2019 | 2020 | 2021 | 2022 | 2023 | 2024<sup>c</sup> |
| --- | --- | --- | --- | --- | --- | --- | --- |
| COA | 0.83 | 0.75 | 0.79 | 0.82 | 0.86 | 0.91 | 0.87 |
| OLY | 0.78 | 0.85 | 0.84 | 0.87 | 0.85 | 0.92 | 0.97 |
| KLA |  | 0.00 | 0.00 | 0.01 | 0.00 | 0.01 | 0.01 |
| CLE |  |  | 0.00 | 0.00 | 0.00 | 0.00 | 0.00 |
| TYE |  |  |  | 0.15 | 0.10 | 0.05 | 0.07 |
| HJA |  |  |  | 0.00 | 0.00 | 0.00 | 0.00<sup>b</sup> |
| CAS |  |  |  | 0.00 | 0.00 | 0.00 | 0.00<sup>b</sup> |
| NWC |  |  |  | 0.00 | 0.00 | 0.00 | 0.00 |
| MAR |  |  |  | 0.00 | 0.00 | 0.00 | 0.00 |
| RAI |  |  |  | 0.09 | 0.13 | 0.13 | 0.14 |
| WA 2% |  |  |  |  | 0.00 | 0.13 | 0.05 |
| OR 2% |  |  |  |  | 0.08 | 0.20 | 0.19 |
| CA 2% |  |  |  |  |  | 0.01 | 0.02 |

<sup>a</sup> In 2022, only Gifford Pinchot National Forest in Washington and Umpqua National Forest in Oregon were sampled at 2%.
<sup>b</sup> No sites reviewed for marbled murrelet.
<sup>c</sup> Only hexagons within the marbled murrelet Northwest Forest Plan conservation zones were reviewed in 2024.

Of the other owl species, northern saw-whet owls and northern pygmy owls (*Glaucidium californicum*) had the greatest number of predicted vocalizations throughout our study (Table 4). We found >1,000 predicted detections of great gray owls in CAS, HJA, and KLA areas (Table 4). Flammulated owls (*Psiloscops flammeolus*) were most predicted (>80,000) in CLE, followed by the 2% sample areas in all three states and NWC (Table 4). Great horned owl and western screech-owl (*Megascops kennicottii*) calls were predicted in all sampling areas (range: 110–86,683 and 1–285,127, respectively). We found a few long-eared owl (*Asio otus*) calls in CAS, CLE, NWC, and 2% areas of all three states (Table 4).

American goshawk, a new sound class in PNW-Cnet v5, were classified at the 0.95 model threshold in all areas; with >4,000 predicted in CAS and >1,000 in CLE and 2% areas of each state. American pikas (*Ochotona princeps*) were commonly detected (>7,000) in CAS, CLE, and WA 2%. Band-tailed pigeon (*Patagioenas fasciata*) call classes were found in all areas with as few as 876 in CLE up to 614,219 apparent detections in COA (Table 4). Sooty grouse (*Dendragapus fuliginosus*) call classes were predicted most densely at OLY (964,279) and TYE (803,328). We reviewed four apparent wolf detections and confirmed one true positive in CLE from July 2024. We forwarded the detection and audio to the Washington Department of Fish and Wildlife Wolf Specialists who confirmed our identification and informed us that the detection overlapped the Naneum pack territory.

We did not find apparent detections for elk (*Cervus canadensis*), ruffed grouse (*Bonasa umbellus*), red-tailed hawk (*Buteo jamaicensis*), bald eagle (*Haliaeetus leucocephalus*), osprey, Wilson’s warbler (*Cardellina pusilla*), evening grosbeak (*Hesperiphona vespertina*), orange-crowned warbler (*Leiothlypis celata*), black-headed grosbeak (*Pheucticus melanocephalus*), chipping sparrow (*Spizella passerine*), pine siskin (*Spinus pinus*), house wren (*Troglodytes aedon*), western flycatcher (*Empidonax difficilis*), Williamson’s sapsucker (*Sphyrapicus thyroideus*), wildcat (*Puma concolor* and *Lynx rufus*), or white-crowned sparrow (*Zonotrichia leucophrys*). Each of these species were new classes for PNW-Cnet.v5 with small training datasets (Lesmeister et al. 2025b), so the lack of apparent detections is likely from poor model performance rather than absence from areas surveyed. For many of these species the recall in PNW-Cnet v5 may have been too low for any true positives to reach the 0.95 classification score threshold used in this report.

## 8. Discussion

We report data collected and summaries for passive acoustic monitoring from 2018–2024 within the NWFP area with emphasis on 2024, the second year for full implementation of the NWFP PAM program using the range-wide 2+20% sampling design (Lesmeister et al. 2021, Lesmeister and Jenkins 2022). Our goal was to survey approximately 1,100 hexagons (4,400 sampling stations) and were able to deploy ARUs within 1,020 hexagons, achieving slightly more than 2023 (Lesmeister et al. 2024), nearly 2.25 million h of recordings processed and apparent detection encounter histories for 68 species. Our efforts provide further evidence that the PAM program has transitioned to a fully operational and analytically mature monitoring system capable of providing range-wide inference on many species’ occupancy, landscape use, and habitat relationships. The integration of 2024 data with recently published, in-review, and in-preparation studies provides a robust framework for understanding species dynamics across the NWFP area and for guiding adaptive management under a changing policy landscape.

The NWFP PAM program was designed to ensure effectiveness for tracking trends in northern spotted owl populations (Lesmeister et al. 2021). By using a random selection of 5-km^2^ hexagons with multiple sampling stations, and collecting data throughout the 24-h period, we have a design suitable for detecting and studying many other forest wildlife species (Tosa et al. 2021, Lesmeister and Jenkins 2022). For example, Rugg et al. (2023) used data collected in 2021 and PNW-Cnet output to evaluate western screech-owl occupancy and the effect of barred owls. Using our data from two study areas, Duarte et al. (2024) demonstrated that the combination of our PAM data and PNW-Cnet are effective to estimate marbled murrelet intensity of use across broad scales and will significantly enhance long-term population monitoring. Using a subset of our PAM data, Weldy et al. (2024) annotated dawn chorus recordings for an open-access dataset that is a valuable resource for researchers developing automated identification tools and those studying the relationship of dawn chorus species to the forest environment.

The primary challenges for the 2024 data collection season were logistical constraints (e.g., landslides, down trees, active logging), early season (March–April) weather with low-elevation snow, wildfires in July and August, and general challenges associated with expanding work into areas unfamiliar to field crews. We continue to gain experience with accessing survey sites and seek continual improvement and efficiencies in the face of significant shortfalls in staffing to support the monitoring program. Once the data are collected, the most significant constraint to producing occupancy estimates for the northern spotted owl continues to be our ability to identify calls originating from call-back surveys conducted by technicians working on other spotted owl projects. These call-back surveys have been the primary method of determining northern spotted owl occupancy for the last 40 years and are widely used by private, state, tribal, and federal entities. The use of a three-note survey tone (USFWS 2021) by some northern spotted owl call-back surveyors has greatly enhanced our ability to screen and identify call-back surveys during data processing with human review for those overlapping survey areas.

In 2024, we expanded partnership networks by collaborating with state and federal agencies. We continue to actively seek partnerships to expand the monitoring network with no additional cost to the program. Additionally, data processing tools developed by the program are freely available and we (PNW Bioacoustics Lab) hosted workshops in 2022–2024 to train state and federal biologists to use the tools for project-level surveys.

### Northern spotted owl

Northern spotted owl occupancy generally increases from north to south (Lesmeister et al. *in review*) and west to east in Oregon. Across the 2021–2024 period, range-wide occupancy estimates remained low, with persistent detections concentrated in the southern Cascades, Klamath Mountains of Oregon and Northwest California. Apparent range contraction in the Oregon Coast Range and Washington Cascades continues, paralleling long-term demographic declines (Franklin et al. 2021, Dugger et al. *in prep*) and spatial patterns of barred owl dominance (Wiens et al. 2021, Jenkins et al. 2025). In 2024, we detected northern spotted owls in 12%, 20%, and 48% of Washington, Oregon, and California hexagons, respectively. For the second year of range-wide surveys, we detected no northern spotted owls north of Stevens Pass in the Washington Cascades sampling area, where 25 of 30 surveyed hexagons had barred owl calling. This may be evidence of extirpation from a region which has had the longest exposure to barred owls. These concordant results across independent datasets (PAM-based occupancy, long-term demographic studies, and invasive species distribution models) provide compelling evidence that barred owl competition remains the primary driver of northern spotted owl displacement throughout most of the NWFP area.

Importantly, the PAM data highlight how occupancy may be stabilizing in southern regions where populations may persist for some time at lower population levels. Together, these patterns suggest that northern spotted owls may persist in limited portions of their historical range where barred owl occupancy remains comparatively low or where landscape configuration provides ecological refugia. This emerging pattern underscores the growing importance of spatially explicit data for identifying potential strongholds and informing management prioritization under the NWFP and federal recovery planning.

### Barred owl

Analyses of 2024 data confirm and expand upon range-wide models of barred owl landscape use in 2023 (Jenkins et al. 2025). The additional year of detections and broader geographic coverage in northern California and the eastern Cascades provides a more complete view of spatial variability in barred owl occupancy. Patterns observed in 2024 were consistent with results from 2023 reported by Jenkins et al. (2025), with the highest rates of landscape use continuing to occur in the Oregon Coast Range and lower rates persisting in the California Coast and Washington Eastern Cascades. This consistency across years reinforces the reliability of the PAM framework for assessing invasive species distributions and offers a robust baseline for evaluating outcomes of targeted barred owl management actions. For example, data generated by our PAM network were instrumental in findings that barred owl management is beneficial for other owl species in addition to spotted owls (Wiens et al. 2025). We continue to collaborate and coordinate with researchers and managers engaged in barred owl management in California and Oregon.

### Marbled murrelet

Marbled murrelet monitoring also advanced substantially in 2024 with contributions of the NWFP PAM program, which provides complementary insights into coastal old-forest ecosystems that parallel the marbled murrelet monitoring framework (Lorenz et al. 2021). Range-wide deployments now encompass over 200 coastal sites from northern California to Washington’s Olympic Peninsula. Analyses of the 2024 NWFP PAM data expanded on the findings of Thomas et al. (2025), which demonstrated that PAM-based murrelet detections closely matched those from traditional audio-visual surveys while reducing costs by more than 70%. Detection probabilities exceeded 0.95 within three weeks of sampling, confirming that PAM is effective for assessing occupancy across extensive, remote coastal terrain (Duarte et al. 2024, Thomas et al. 2025).

Naïve occupancy rates of marbled murrelets in the COA and OLY study areas have remained mostly consistent since 2018, and TYE and RAI since 2021. Naïve occupancy rates for areas within the expected range were 52% overall, 62% in Washington, 57% in Oregon, and 3% in California. Patterns are strongly associated with mature coastal conifer forests and reduced edge disturbance, consistent with established nesting habitat relationships (Raphael et al. 2018, Lorenz et al. 2021). These findings confirm that PAM methods can reliably contribute to efforts to track population use of inland nesting habitat. Together, these results position the NWFP PAM program as a central component of marbled murrelet population monitoring, providing a scalable approach to support adaptive management of coastal forests under the NWFP.

### Biodiversity

Beyond northern spotted owl and marbled murrelet monitoring, the NWFP PAM program now supports a growing portfolio of biodiversity monitoring applications across the NWFP region. Autonomous recording units deployed under the 2+20% sampling design detect a wide array of taxa, including forest songbirds, bats, amphibians, and small mammals, in addition to tracking anthropogenic and geophysical sound sources. These “bioacoustic bycatch” detections provide a unique window into ecosystem function, species interactions and distribution, and disturbance recovery processes that extend far beyond individual target species.

Several analyses have expanded beyond core species to quantify seasonal and community-level acoustic diversity metrics across major ecological provinces. For example, Habib et al. (2025) found that sensory interference plays an important role in northern saw-whet owl habitat. Further, preliminary results indicate that soundscape diversity remains highest in late-successional forests and in areas with moderate fire frequency, suggesting that structural heterogeneity and fire refugia support higher overall biodiversity and occupancy of forest-adapted species (e.g., Ruff et al. *in press,* Kohlberg et al. *in review*). These findings align with the broader ecological literature linking compositional complexity to resilience and provide an empirical framework for evaluating ecosystem conditions under the NWFP.

By integrating soundscape metrics with spatial data on vegetation, fire, and climate the NWFP PAM program can evolve into a multi-taxa biodiversity observatory capable of addressing core Forest Service and partner agency objectives for disturbance adaptation, ecosystem resilience, and sustainable resource management. This capability strengthens the position NWFP effectiveness monitoring as one of the most comprehensive, landscape-scale biodiversity monitoring efforts worldwide by linking population dynamics, habitat change, and ecosystem processes through a single, standardized data stream (Lesmeister and Jenkins 2022).

The 2024 analyses also reflect substantial progress toward the integration of species, habitat, and disturbance monitoring components envisioned for biodiversity monitoring by the NWFP Effectiveness Monitoring Program (Mulder et al. 1999). Following the framework described by Lesmeister et al. (*in prep*), we suggest that the analytical workflow described here could be expanded to more directly incorporate spatial data on forest structure and disturbance history (Davis et al. 2022b) and fire refugia to evaluate how these factors influence species occupancy and resolve conflicts between conservation and management (Lesmeister et al. 2025a). Preliminary models indicate that northern spotted owl occupancy remains highest in areas with greater proportions of mature and late-successional forests and lower in areas with recent disturbance. Areas with repeated low- to moderate-severity fire continue to support consistent detections of spotted owls, aligning with broader findings linking postfire forest resilience to owl persistence (Rockweit et al. 2024).

These results underscore the analytical advantages of integrating PAM-based occupancy data with landscape-scale habitat indicators. Unlike prior demographic monitoring, which was spatially limited to fixed study areas, the PAM framework supports range-wide inference with explicit connections to habitat metrics derived from remote sensing and forest inventory data (Lesmeister and Jenkins 2022). This integration enhances the capacity of NWFP Effectiveness Monitoring to assess both species-specific responses and broader ecosystem resilience to disturbance and early signals of disturbance. For example, we have several human disturbance noise classes in PNW-Cnet v5 that can be used to quantify changes in human activity in Pacific Northwest forests. In 2024, the Oregon study areas with a checkerboard pattern of private and federal ownership (COA, KLA, TYE, and the OR 2%) have far more logging yarding system whistle (i.e., yarder machine) detections (222,569) compared to other regions (14,440), and is more than four times the number of yarders we detected in 2023 (Lesmeister et al. 2024). We expect that our PAM data will be increasingly informative with long-term monitoring to quantify occupancy dynamics in relation to these disturbance indicators. We will continually seek to generate new insights into these ecosystems due to the flexibility and scale of the biodiversity monitoring program.

### Outlook

The collective findings from 2024 reinforce that northern spotted owl populations remain in steep decline and that barred owl competition continues to be the dominant limiting factor across the range. However, the convergence of demographic, acoustic, and habitat data also reveals key areas of persistence that provide hope for targeted conservation action. The emerging integration of PAM with spatial habitat and disturbance indicators now enables the NWFP monitoring program to move beyond descriptive status assessments toward predictive and adaptive modeling that can directly inform management.

Looking ahead, maintaining and expanding the PAM network will be critical for sustaining this analytical capacity. Continued cross-agency coordination will ensure that data collection, processing, and analysis remain standardized across years, enabling consistent trend estimation and integration with complementary datasets such as forest inventory, remote sensing, and fire monitoring. The PAM program’s demonstrated capacity to produce peer-reviewed scientific outputs (e.g., Appel et al. 2023, Rugg et al. 2023, Weldy et al. 2023, Duarte et al. 2024, Habib et al. 2025, Jenkins et al. 2025, Rugg et al. 2025, Ruff et al. *in press*) attests to its scientific rigor and relevance to informed management. Together, these efforts establish the foundation for a next-generation monitoring program that links species and ecosystem processes across space and time, supports adaptive management and provides a model for large-scale biodiversity monitoring (Tosa et al. 2021).

At the same time, sustaining this capacity depends on retaining the scientific and technical workforce that built it. The loss of field biologists, data analysts, and technical specialists through program cuts and attrition poses a serious risk to long-term continuity. The expertise embedded in these teams that has been lost in recent cuts includes working in remote mountainous forests, acoustic data management, species identification, analytical modeling, and forest ecology. A contraction in field staffing would reduce the program’s ability to maintain the PAM network, degrade data quality, and limit responsiveness to emerging management needs.

The implications of reduced sampling effort are well-documented. Simulation analyses conducted by Lesmeister et al. (2021) demonstrated that smaller annual sample sizes substantially decrease confidence in occupancy estimates and reduce statistical power to detect biologically meaningful changes in species status at both local- and range-wide scales. These findings highlight the quantitative risk of future program reductions: diminished field capacity would directly translate into weaker inference, eroding the scientific credibility of long-term trend estimates. It would also undermine the Forest Service’s ability to provide defensible, science-based information necessary for balanced timber harvest planning under the NWFP.

The PAM program has already demonstrated its potential to support many timber projects, identify harvest-compatible zones, quantify habitat condition, and evaluate post-treatment effects on wildlife occupancy. Eroding the human and analytical infrastructure that supports these analyses would therefore diminish both conservation and management outcomes. Protecting field and analytical capacity is essential to maintaining the integrity of NWFP monitoring, safeguarding its role in informing sustainable forest management, and preserving one of the few range-wide, science-based monitoring systems capable of addressing biodiversity and resource management challenges at landscape scales.

Funding and support for this program was provided by: USDI Bureau of Land Management and National Park Service; USDA Forest Service Pacific Northwest Region and Pacific Northwest Research Station. We thank our partners at Oregon State University, C. Appel, M. Weldy, A. Kohlberg, T. Levi, C. Sullivan, J. Koning, B. Padmaraju, M. Samaduroff, and A. Subramanian for assistance in PNW-Cnet development and data processing. We are deeply indebted to the dedicated field and lab technicians that collected and processed millions of hours of bioacoustics data: M. Alsaid, G. Alvarez, M. Barnett, H. Basile, E. Bernier, A. Brown, A. Carlson, I. Comcowich, J. Crawford, S. Diaz, J. Domeika, T. Ellis, J. Fitzgerald, A. Groszek, E. Guzman, A. Habib, M. Holmgren, A. Ingraham, D. Jackson, K. Kadoun, K. Kimball, B. Kintner, A. Koehler, P. Loafman, J. Martineau, S. Marxer, N. McClain, D. Medeiros, S. Mirotznik, T. Munger, M. O’Neill, E. Ostraff, K. Parker, R. Pechtimaldjian, W. Perkins, B. Plumhoff, L. Potvin, C. Provost, K. Rider, X. Rongel, D. Schapker-Mendez, C. Sendek, C. Silva, K. Skaug, M. Smith, B. Soto, D. Starzenski, C. Stephens, C. Urnes, E. Yargeau, N. Younan, Olympic Wildland Fire Module, Baker River Hotshots, and all other crew members, collaborators, and volunteers who assisted on the project in some capacity.

